# When and why grid cells appear or not in trained path integrators

**DOI:** 10.1101/2022.11.14.516537

**Authors:** Ben Sorscher, Gabriel C. Mel, Aran Nayebi, Lisa Giocomo, Daniel Yamins, Surya Ganguli

**Affiliations:** Department of Applied Physics, Stanford University; Neurosciences PhD Program, Stanford University; McGovern Institute for Brain Research, MIT; Department of Neurobiology, Stanford University School of Medicine; Department of Psychology, Stanford University; Department of Computer Science, Stanford University

## Abstract

Recent work has claimed that the emergence of grid cells from trained path-integrator circuits is a more fragile phenomenon than previously reported. In this note we critically assess the main analysis and simulation results underlying this claim, within the proper context of previously published theoretical work. Our assessment reveals that the emergence of grid cells is entirely consistent with this prior theory: hexagonal grid cells robustly emerge precisely when prior theory predicts they should, and don’t when prior theory predicts they should not.

## 1 Introduction

Recent works [1, 2, 3, 4] have found that training neural circuits to path integrate (i.e. convert velocity inputs into a desired place cell output code through a set of hidden units) can lead to the emergence of grid-like units [5, 6] with square [1] heterogeneous [2] or hexagonal [3, 4] lattice structures, under some simple biologically plausible constraints. This raises interesting theoretical questions about when and why grid cells emerge from trained path-integrators and what their lattice structure might be (square, hexagonal, or heterogeneous). These questions were addressed in [3, 4].

Subsequent work [7] presented a sequence of simulation results and some theoretical analysis suggesting prior work involved non-transparent fine-tuning. In this note we critically assess the simulations and theory presented in [7]. Our assessment leads to the following outline and main conclusions for this note:

1. Since [7] did not explain prior theory accounting for when grid cells emerge or not from trained path-integrator circuits, we begin in Sec. 2 by reviewing the theory of [3, 4], especially as many results claimed to be surprising in [7] actually follow from the theory in [3, 4].
2. [7] claims that finding hexagonal grids in trained path-integrators is an extremely fragile phenomenon because training 11,000 path-integrators resulted in hexagonal grid cells only 10% of the time. Based on this [7] suggested that prior work involved non-transparent fine tuning. In Sec. 3 we discuss how this claim is misleading, because many choices made in [7] were previously shown by theory *not* to lead to hexagonal grid cells. Indeed when [7] conditioned on two key properties shown in prior theory [3, 4] to lead to the emergence of hexagonal grid cells, [7] itself robustly found hexagonal grid cells close to 100% of the time without any fine-tuning.
3. A main claim in [7] is that a highly specific place cell output code is required to generate hexagonal grid cells, namely a difference of softmaxes (DoS) structure, and that other proposed structures like differences of Gaussians (DoG) do not suffice. We show in Sec. 4 that this claim is incorrect, due to a normalization error in [7], and that DoG place cell input structure can reliably generate hexagonal grids as predicted by theory in [3, 4].
4. [7] claimed that place cells with multiple fields cannot generate hexagonal grid cells. We show in Sec. 5 that place cells dominated by a single scale with multiple fields can indeed generate hexagonal grid cells, consistent with the theory given in [3, 4]. We further discuss the role of multiple scales.
5. [7] claims that extremely small changes in place scale can lead to complete disappearance of grid cells (e.g. 10cm and 12cm place cell scales can generate grids but 11cm cannot). We show in Sec. 6 that this is not the case in trained path-integrators that robustly generate a large fraction of hexagonal grid cells with high grid score. The networks studied in [7] do not generate grid cells as robustly, suggesting that the variations in grid score distributions with respect to place cell scale observed in [7] may simply reflect statistical noise in path-integrators that do not robustly exhibit grid cells to begin with.
6. [7] presents a proof that Gaussian place cells cannot generate periodically patterned grid cells in trained path integrators. We show in Sec. 7 that the conclusion of this proof is inconsistent with simulations. We further resolve the discrepancy between the claimed proof in [7] and simulations by showing how the proof in [7] is incomplete and how the theory in [3, 4] suggests periodic patterns are indeed possible with Gaussian place cells.
7. [7] claims that DoG/DoS structure in place cells is biologically unrealistic. We discuss in Sec. 8 that when adhering to the precise interpretation of DoG/DoS structure, given in [4], as that of grid cell inputs to place cells, rather than the rectified outputs of place cells, such DoG/DoS structure is not obviously implausible given the current lack of the requisite experiments to rule it out.
8. We summmarize our critical assessment of the claims made in [7] in Sec. 9.

### 2 Review of theory for the emergence of hexagonal grid cells

We begin by giving a concise review of theory in [3, 4] explaining when grid cells appear or not in trained path-integrators, as this theory is helpful for assessing claims made in [7]. Overall, our theory provides a unifying conceptual explanation for when and why square, heterogeneous or hexagonal grids spontaneously emerge in trained path-integrators. The main conclusion is that trained path-integrators that convert velocity inputs into desired place cell input currents through a single layer of hidden units with minimal possible activity, will learn grid cell patterns in the hidden units as long as the place cell currents have a relatively narrow center-surround structure and the hidden units are constrained to have nonnegative firing rates. A reader who is willing to take this high level conclusion on faith can skip this section.

A detailed derivation of the theory is given in [4]; here we simply describe the mathematical conclusions of the theory. We begin by noting that any network that converts velocity inputs to place cell representations through a set of hidden units must solve a position encoding problem, schematized as 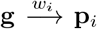. Here **g** denotes an n_*x*_ dimensional firing rate vector of a single hidden unit in the network across n_*x*_ spatial bins.

When the animal is at spatial bin *x* this cell’s firing rate is given by g(*x*). Similarly **p**_*i*_ for i = 1, …, n_*p*_ denotes the firing rate vectors of n_*p*_ place cells with similarly defined rate maps p_*i*_(*x*). More precisely, in our interpretation, p_*i*_(*x*) denotes the *input currents* to place cell i when the animal is at location *x, before* they are rectified to generate positive place cell output firing rates.

We show in [3, 4] that the nature of the representations learned by the hidden units can be gleaned by solving the auxiliary position encoding problem

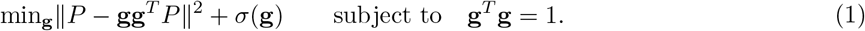

Here, we have collected the place cell input current vectors as the n_*p*_ columns of the n_*x*_ by n_*p*_ matrix P. The first term forces the hidden unit (i.e. putative grid cell) to accurately generate place cell input currents, while the second term is a negativity cost *σ*(g) that is large (small) if g is negative (positive). Minimizing the negativity cost promotes positive firing rates in the putative grid cell. The solutions to the position encoding optimization problem in (1) can be understood in terms of the correlational structure of the place cell input currents. Indeed the correlational structure of outputs in a learning problem often determine hidden unit representations, as seen in both the development of semantic categories [8, 9] and the development of ocular dominance columns [10]. In our grid cell case, the correlational structure of desired place cell input currents is described by the n_*x*_ by n_*x*_ matrix with matrix elements

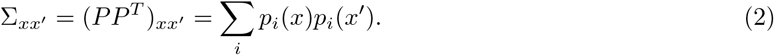

Here a positive (negative) matrix element Σ_*xx′*_ quantifies how similar (dissimilar) the place cell input current vector is at two points *x* and *x*′ in space. Figure 1A shows a single row of the spatial correlation matrix where *x* is the center of a 2D enclosure, and *x*′ varies over the enclosure, in the case where the place cell input currents p_*i*_(*x*) have a center-surround structure.

**Figure 1:**
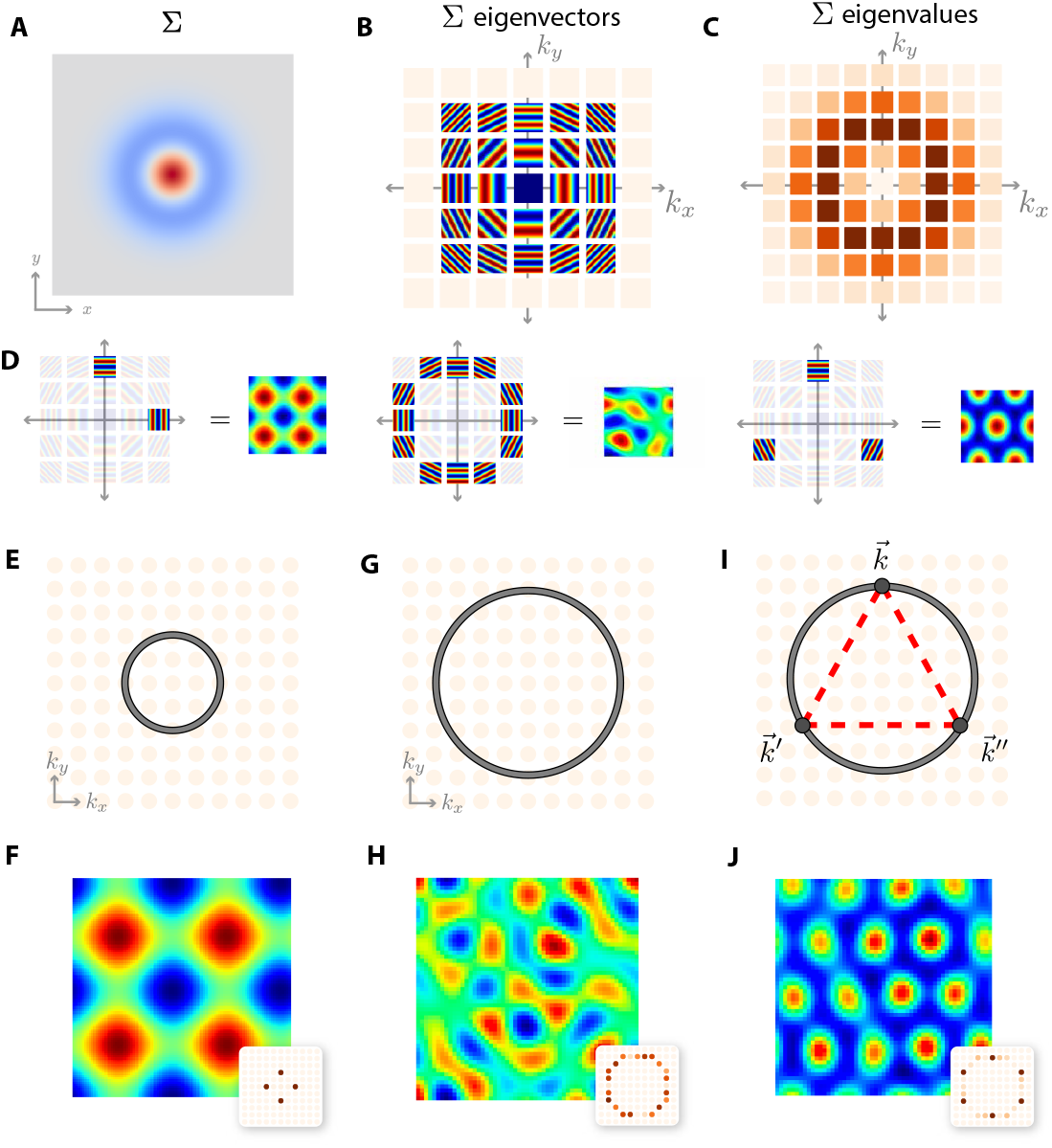
A theory for predicting the structure of learned spatial maps. (A) A central row of the place cell correlation matrix Σ, indicating similarity in the target place cell code as a function of spatial displacement, illustrating center surround structure. Red (blue) indicates positive (negative) similarity. (B) Eigenvectors of Σ are well approximated by Fourier plane waves, and are shown arranged on a discrete lattice of integers (k_*x*_, k_*y*_) corresponding to the frequencies of each plane wave along the two cardinal axes. **(C)** The eigenvalues of Σ corresponding to the eigenvectors in B. Darker red indicates larger eigenvalues. **(D)** Particular linear combinations of plane waves yield grid patterns. A square grid can be generated by combining waves that oscillate along the cardinal axes (left). A heterogeneous grid can be generated by arbitrary combinations of waves with frequencies near an annulus of fixed radius (middle). A hexagonal grid pattern arises when waves whose frequencies form an equilateral triangle in the (k_*x*_, k_*y*_) plane are combined (right). (E) If the similarity structure in panel A is wide, the annulus of Σ eigenvalues will be small, shown in grey, and will only intersects a few plane modes oscillating primarily along the cardinal axes. (F) Simulations confirm that neural circuits with unconstrained firing rates will learn combinations of precisely these cardinal modes, generating square grids. (G) For narrow similarity structure, the large eigenvalues of Σ lie near an annulus of large radius, shown in grey. (H) Simulations confirm that neural circuits with unconstrained firing rates learn arbitrary linear combinations of plane waves with oscillations frequencies near this annulus, generating heterogeneous grids. (I) The nonnegativity constraint creates a cooperative interaction among frequency triples that sum to 0 as vectors in the (k_*x*_, k_*y*_) lattice. Since these frequencies must lie on the annulus of large eigenvalues, they must form an equilateral triangle (dashed red). (J) Simulations confirm that neural circuits with nonnegative firing rates learn hexagonal grids, as predicted in panels D (right) and I. Panels F,H,J show the learned grid cell representation and Fourier power spectrum (inset).

Now the importance of Σ lies in the fact that without the negativity cost *σ* in (1), an optimal solution corresponds simply to the top principal eigenvector of Σ with largest eigenvalue. When there are many eigenvectors with the same top degenerate eigenvalue then there can be many optimal solutions to the position encoding problem consisting of arbitrary linear combinations of these top eigenvectors. Therefore we need to understand the eigenstructure of Σ which will play a *key role* in the nature of learned grid cells. Assuming translation invariant statistics of place cell input currents (which is approximately true up up to boundaries), the eigenvectors of Σ are well approximated by Fourier plane waves that oscillate in different frequencies and directions (Figure 1B). Thus the eigenvectors are indexed by two integers, k_*x*_ and k_*y*_, indicating the spatial frequency of oscillation in each of the two cardinal spatial directions. Each such eigenvector has an associated nonnegative eigenvalue. These eigenvalues are shown in Figure 1C at the integer (k_*x*_, k_*y*_) lattice points associated with the corresponding eigenvectors in Figure 1B. The strength of these eigenvalues can be obtained by computing the power in each Fourier mode at spatial frequency (k_*x*_, k_*y*_) of the similarity function displayed Figure 1A. Because this similarity function has a center-surround structure, the maximal eigenvalues, or equivalently Fourier power, occurs near an annulus in the (k_*x*_, k_*y*_) lattice, and the narrower the similarity function in Figure 1A, the larger the radius of this annulus in Figure 1C. Indeed hexagonal grid cells were robustly found in [3, 4] in trained path-integrators based on this theory.

Crucially, as Figure 1C shows, there are multiple maximal eigenvalues distributed over a ring centered on the origin, corresponding to multiple top eigenvalues of Σ. Consequently, without the negativity cost *σ*, there is an entire family of equally optimal grid maps consisting of arbitrary linear combinations of eigenvectors in Figure 1B with maximal associated eigenvalue. Figure 1D indicates how, for example, square, heterogeneous, or hexagonal grid patterns can be constructed from appropriate combinations of these top eigenvectors. This allows us to answer how the structure of the place cell input current vectors interact with the presence or absence of the negativity cost *σ*(**g**) to generate these three types of grid codes.

In the first case, if the place cell input similarity structure in Figure 1A is wide relative to the size of the enclosure, then the maximal eigenvalues will occur near an annulus of small radius, as in Figure 1E. This annulus will intersect a small number of lattice points in the (k_*x*_, k_*y*_) plane corresponding to low frequency eigenmode oscillations aligned along the cardinal axes of the enclosure, and linear combinations of these oscillations along these cardinal directions would predict square grid cells as in Figure 1D, left. This prediction is confirmed in simulations of our position encoding problem in Figure 1F. Indeed, square grid cells were previously found in trained path integrators [1] where they used extremely low frequency desired place cell outputs (indeed linear). On the other hand, if the similarity structure in Figure 1A is narrow, the maximal eigenvalues will occur near an annulus of large radius, which intersects many lattice points in the (k_*x*_, k_*y*_) plane, as in Figure 1G. As described above, without further constraints *σ*(**g**), the hidden representation of neural circuits will learn arbitrary linear combinations of eigenvectors associated with the many lattice points on the large annulus, yielding relatively heterogeneous patterns, as predicted in Figure 1D, middle. This prediction is confirmed in simulations in Figure 1H. Indeed [2], with very narrow place cell tuning, found highly heterogeneous grid-like representations, with a few cells having a high grid score, but the entire distribution of grid scores was indistinguishable from that obtained by grid-patterns obtained by low-pass filtering random noise as demonstrated in [4].

We now turn to the effect of the negativity cost *σ*(**g**), whose minimization must break the tie amongst all equally optimal maps consisting of linear combinations of all eigenvectors found on the ring of maximal eigenvalues in Figure 1B,C. We proved in [4] that the effect of this nonnegativity cost is to favor 3-fold combinations of eigenvectors whose spatial frequencies (k_*x*_, k_*y*_) form an equilateral triangle centered at the origin (Figure 1I). This combination of eigenvectors predicts a hexagonal grid representation as in Figure 1D, right. This prediction is confirmed in simulations of our position encoding problem in Figure 1J.

### 3 Hexagonal grid cells robustly emerge from trained path-integrators under two key conditions predicted by prior theory

A main take home message in [7] is that training 11, 000 path-integrator networks under various choices lead to hexagonal grid cells less than 10% of the time, implying that prior work involved non-transparent fine-tuning. Indeed [7] claimed that its search over various choices “*was biased toward configurations shown to produce grid cell emergence and thus our findings about the fragility of these solutions conservatively favored these solutions as much as possible*.” This claim is incorrect. Indeed only a small fraction of the many choices made in [7] were shown by prior theory in [3, 4] to lead to hexagonal grid cells. For example, only 2 out of the 4 nonlinearities examined in [7] (ReLU, Sigmoid) have a nonzero negativity cost shown in Sec. 2 to favor hexagonal grids. Also only 1 out of the 5 desired place cell structures is consistent with the Fourier annulus spectral structure of place cell correlations shown schematically in Fig. 1C. In the experiments [7] performed which included these two ingredients, [7] actually robustly obtained hexagonal grid cells close to 100% of the time (in [7] see Fig. 2A bottom row brown bar corresponding to difference of softmax (DoS)). Indeed, the fact that only 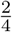 nonlinearities should work and only 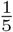 place cell structures should work, means prior theory actually correctly predicts that [7] should only expect hexagonal grid cells 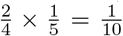 of the time, given their choices. This is indeed roughly the fraction of times [7] sees hexagonal grid cells across all choices made.

**Figure 2:**
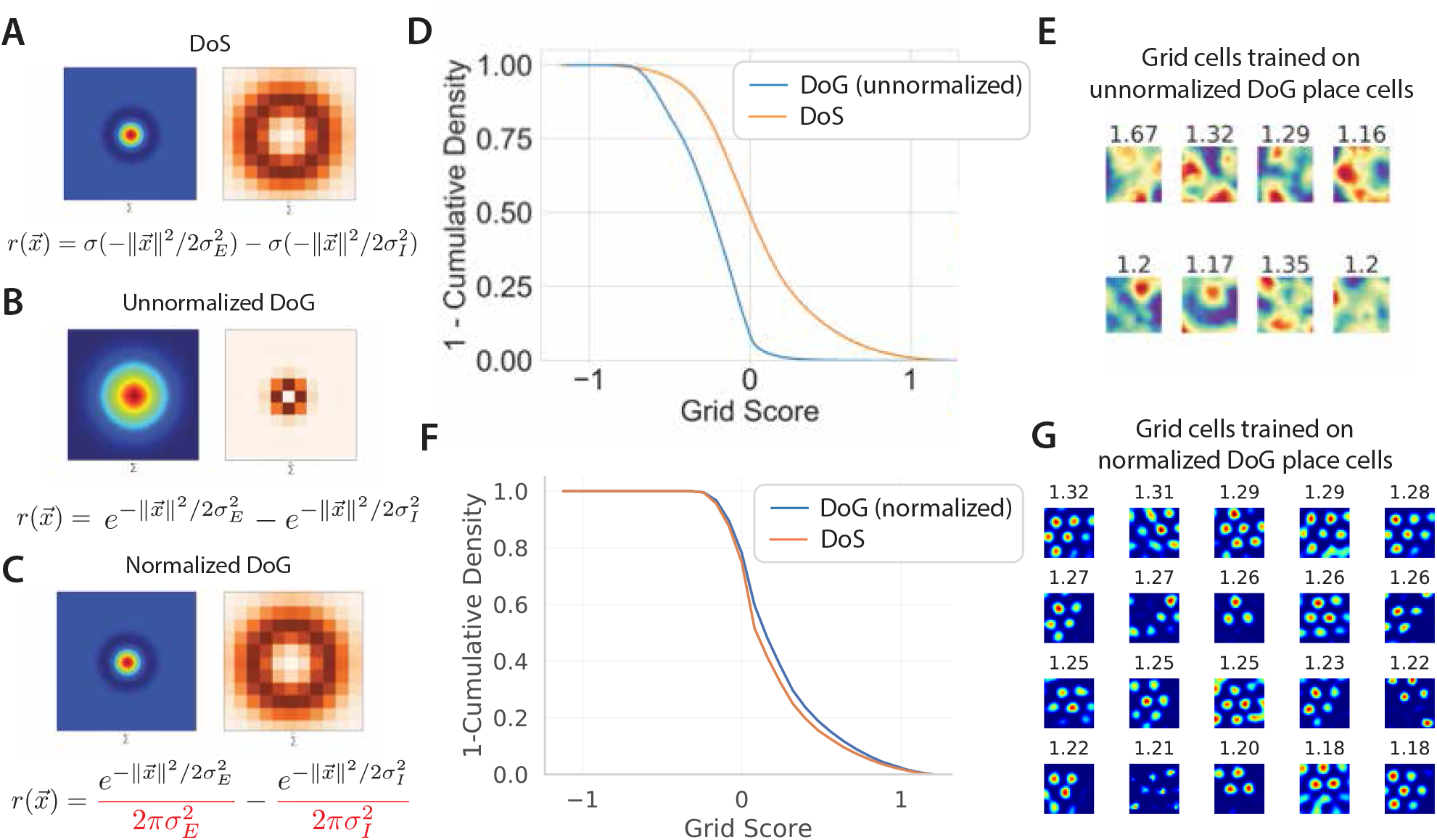
Grid cells emerge equally reliably for DoS and DoG place cell input structures. (A,B,C) Place cell correlational structure in the spatial domain (left) and spatial frequency domain (right) for DoS structure (A), improperly normalized DoG structure (B), and properly normalized DoG structure (C). These figures follow conventions used in Fig. 1AC. (D,E) Reproduced from [7]: path integrators trained to produce improperly normalized DoG place cell input structure do not produce hexagaonal grid cells, consistent the theory in [3, 4]. Training with unnormalized DoGs (Eq. 3) leads to a lower distribution of grid scores than DoS (D) and cells that are not grid-like (E). (F,G) Training with properly normalized DoG place cell input structure leads to an identical distribution of grid cells as DoS (F) and reliably produces regular hexagonal grid cells (G). Grid score distribution in (F) are computed over 10 independently trained models each for DoG and DoS, with identical hyperparameters and different random weight initializations, simulated trajectories, and place cell centers. Hyperparameters were chosen to match [7], except that models were trained for 1/10 as many steps with 10x learning rate to speed training. (G) shows the top grid cells from one such trained network for direct comparison with (E).

Thus the initial result of [7], far from revealing non-transparent fine-tuning in prior work, is essentially consistent with known theory in [3, 4]. Indeed hexagonal grid cells robustly emerge in [7] precisely when [3, 4] predicts they should, and otherwise they do not appear precisely when [3, 4] predicts they should not.

### 4 DoG place cell input structures can yield hexagonal grids

[7] claims that a very specific Difference of Softmaxes (DoS) place cell structure is required for the emergence of hexagonal grid cells, and a Difference of Gaussian (DoG) structure will not lead to this emergence (see Fig. 4bc in [7]). This claim is incorrect. Hexagonal grid cells emerge equally reliably for *both* DoG and DoS target functions as we demonstrate in Fig. 2. The error in [7] occurs because the Gaussian function was improperly normalized in [7]. The following form for DoG place cells was used ([7], Appendix A):

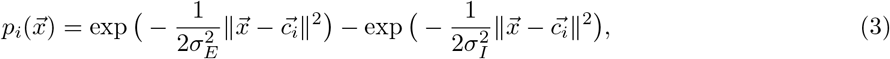

while the properly normalized expression is given by

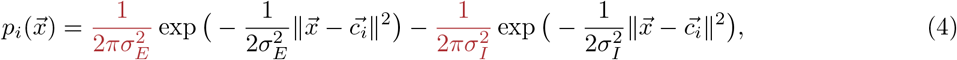

Where 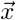 is the animal’s 2d position, 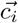 is the place cell receptive field center, and 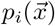 is the place cell input current to cell i when the animal is at position 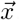. Note the unnormalized DoG does not exhibit a large radius annulus of high Fourier power in the spatial frequency domain (Fig. 2B) but the properly normalized DoG does (Fig. 2C). Then according to theory from [3, 4] reviewed above, the incorrectly normalized DoG function should not be expected to lead to hexagonal grids (Fig.2DE), but the properly normalized DoG should, as we demonstrate in (Fig.2FG). Indeed a key point of the theory in [3, 4] is the necessity of a large radius annulus structure in the Fourier domain for place cell inputs. [7] simply used an incorrect normalization that removes this structure, but as we show, this does not mean that no DoG place cell structure can yield hexagonal grids.

In summary, we have demonstrated, in contrast to the claim made in [7], that DoG place field structure can indeed lead to the emergence of hexagonal grids as long as its correlational structure exhibits a large radius annulus of high power in the Fourier domain, as predicted by theory in [3, 4].

### 5 Place cells with multiple fields can yield hexagonal grid cells

[7] demonstrates that training path-integrators to produce place cells with multiple scales and multiple fields per place cell does not lead to hexagonal grids (see Fig. 5 in [7], also copied in our Fig. 3AB). Assuming that in the Fourier domain this multiscale place field structure leads to a disk of large eigenvalues spanning many spatial frequencies instead of an annulus concentrated at a single radius in the spatial frequency domain, as in Fig. 1C and Fig. 2AC, then the lack of hexagonal grid cells under multiscale, multibump place cell structure is entirely predicted by the theory in [3, 4].

**Figure 3:**
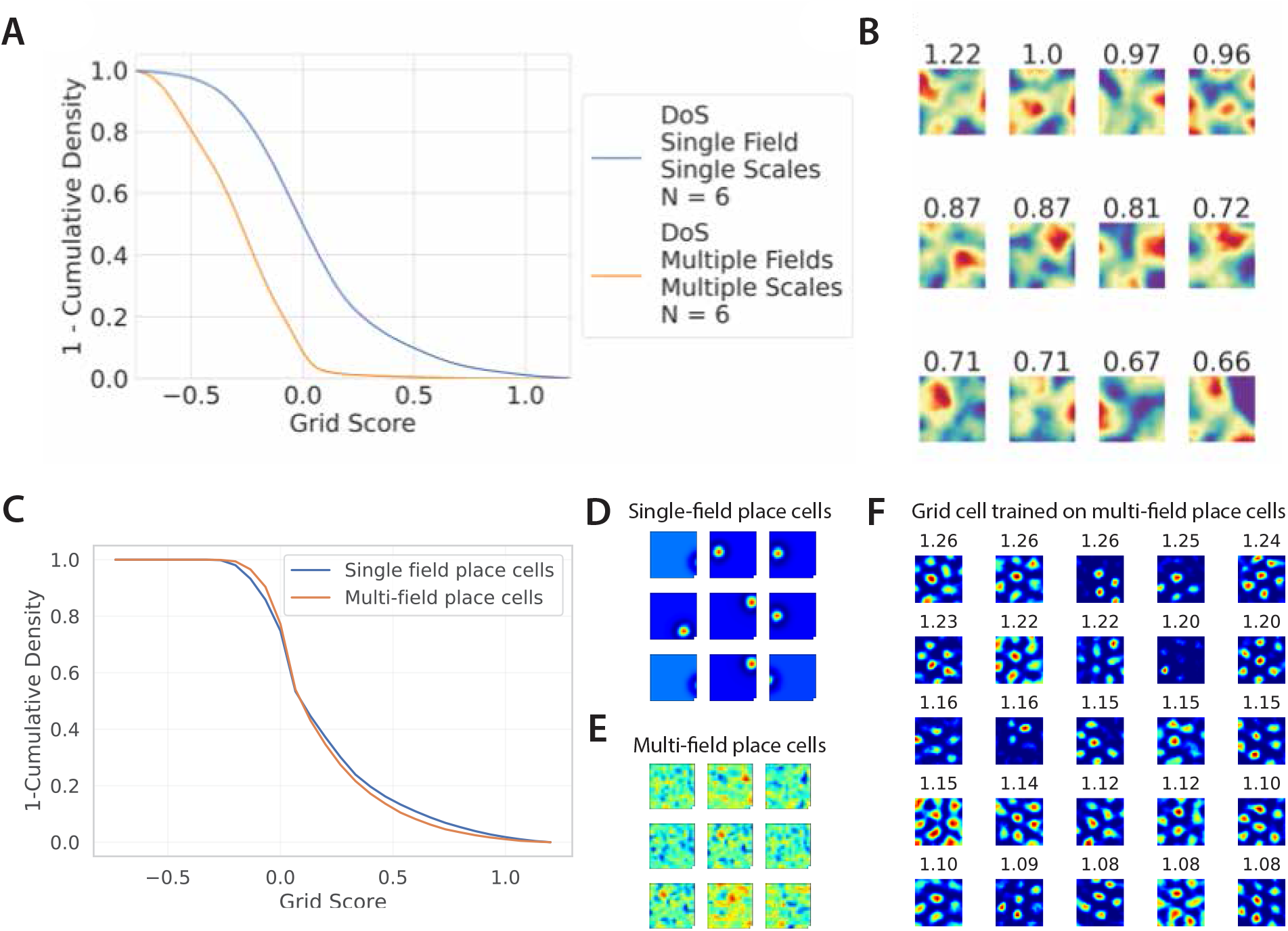
Place cells with multiple fields can yield hexagonal grid cells. The theory in [3, 4] shows that the key determinant of grid cell structure in trained path integrators is not whether the place cells targets have single fields or multiple fields, but whether they have the correct correlation structure. Changes to place cell tuning curves which disrupt this correlation structure should lead to a lower distribution of grid scores. (A,B) Reproduced from [7], show that changing the place cell tuning curves by adding multiple fields and multiple scales, without preserving the correlation structure, can lead to a lower distribution of grid scores (A) and cells that are not grid-like (B). (CDEF) show that changing the place cell tuning curves by adding multiple fields while *preserving* the place cell correlation structure leads to no change in the distribution of grid scores (C). Grid score distribution in (C) is computed over 10 independently trained models using either single-field place cells (D) or multi-field place cells (E) generated as random Gaussian tuning curves with the same 2d correlation structure as the single-field place cells (Fig. 2A). Models are trained with identical hyperparameters and different random weight initializations, simulated trajectories, and place cell centers. Hyperparameters were chosen to match [7], except that models were trained for 1/10 as many steps with 10x learning rate to speed training. (F) shows the top grid cells from one such trained network for direct comparison with (B).

**Figure 4:**
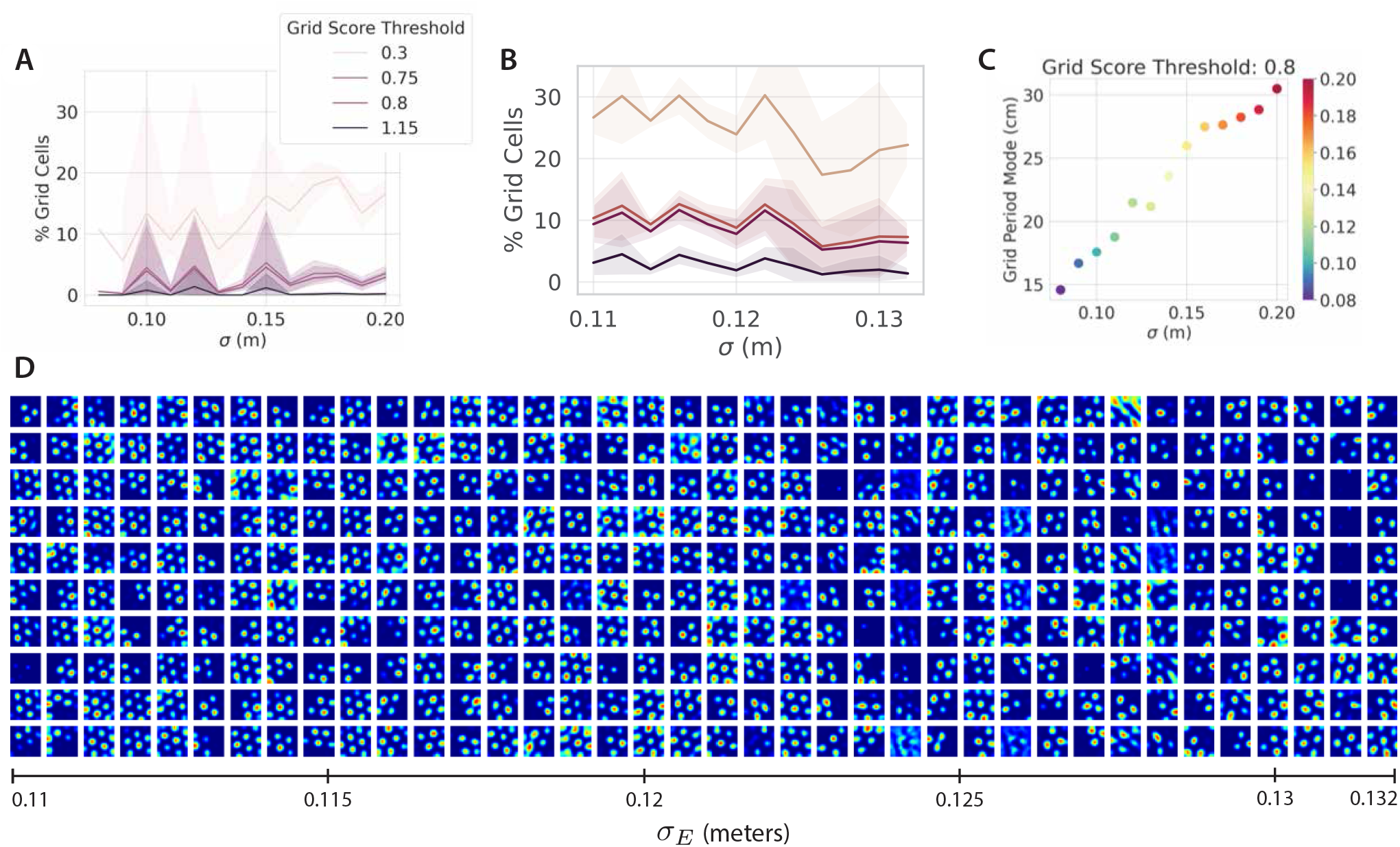
Hexagonal grids are robust to small changes in place cell scale. We empirically investigate the claim in [7] that small changes in the place cell targets’ receptive field width *σ* result in the disappearance of grid cells (A, reproduced from [7]). We trained 120 models, sweeping *σ* over a 20% range. All models were trained with identical hyperparameters and different random weight initializations, simulated trajectories, and place cell centers. (B) We found that all trained networks, for all values of *σ*, produced grid cells with grid score greater than any of the thresholds used in [7]. Moreover the grid cell scale increased smoothly with *σ*, consistent with [7]’s own findings (C, reproduced from [7]). (D) shows the top grid cells from each trained network, sorted by *σ*, clearly demonstrating that each trained network learns regular hexagonal grids which smoothly increase in scale with increasing *σ*. Each column shows the top 10 grid cells from one of 38 randomly chosen trained networks, and the networks are organized along the x-axis by the target receptive field width *σ* they are trained with. Hyperparameters were chosen to match [7], except that models were trained for 1/10 as many steps with 10x learning rate to speed training of a large number of networks.

**Figure 5:**
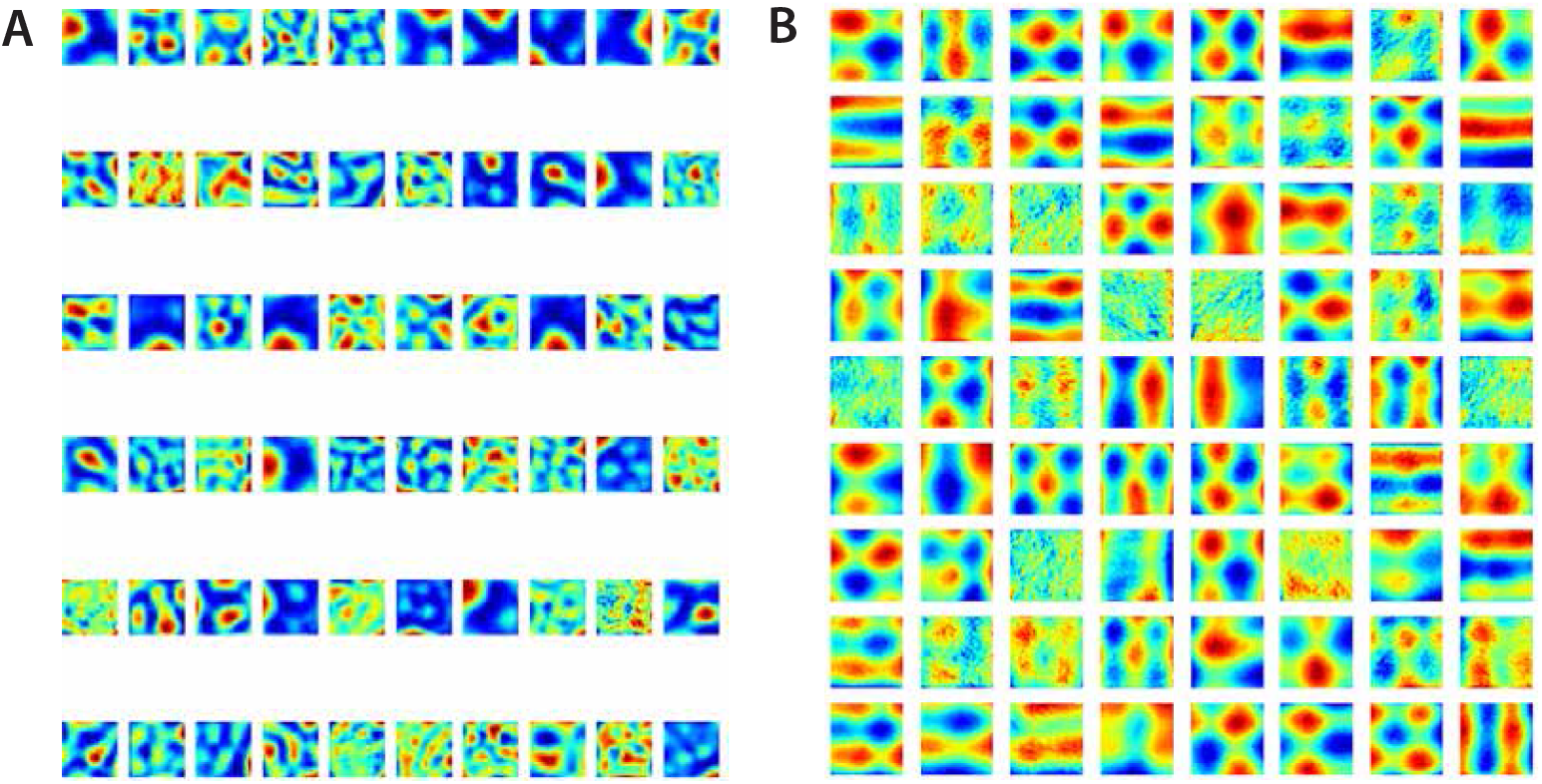
Gaussian place cells can generate periodic patterns. (A) Reproduced from [2]: a path integrator trained on Gaussian place cells targets with width 1cm produces irregular periodic patterns. (B) An adapted version of the code used to generate (A), with Gaussian place cell targets with width 5cm, produces stripes and square periodic patterns. The model and training hyperparameters can be found at https://github.com/ganguli-lab/grid-pattern-formation.

However, place fields of multiple scales are primarily found at different locations along the dorsoventral axis of the hippocampus [11, 12]. Moreover, grid cells of different scales are found at corresponding locations along the dorsoventral axis of the medial entorhinal cortex (MEC) [5, 13]. This raises a natural question: if we restrict our attention to modelling a single location along the dorsoventral axis, corresponding to a single scale for place cells, but we include multiple firing fields for place cells, will hexagonal grid cells naturally appear? As we show, the answer is yes, as already predicted by prior theory.

As explained in [3, 4], a key ingredient for generating hexagonal grid cells is not unimodal place cell input currents or firing fields, or indeed even individual center surround input currents, but rather center-surround *correlational statistics* in the input currents to place fields (i.e the structure of Σ in (2) shown in Fig. 1AC). To demonstrate the primary theoretical importance of this correlational structure, and to demonstrate the complete unimportance of single bump unimodal place cell input structure, we generate completely *random* multiple bump place cell input patterns p_*i*_(x) constrained to have the same annulus like center-surround correlational structure as in Fig. 1AC. As we illustrate in Fig. 3**CDEF**, this leads to highly heterogenous place cell inputs (Fig. 3**E**), that still nevertheless generate clear hexagonal grid cells (Fig. 3**CF**). Thus both theory and experiment demonstrate that unimodal place cell input currents or firing fields are not necessary for the emergence of hexagonal grid cells; heterogenous place cells with a single-scale annulus-like correlational structure can robustly generate hexagonal grid cells, where the scale of the grid cell is determined by the scale of the place cell. All of this was predicted by the theory of [3, 4].

Questions regarding multiple grid scales were outside the scope of [3, 4]. However, as discussed above, the practice of studying MEC phenomena one scale at a time (at least approximately), is motivated by the biological fact that grid scale is organized topographically along MEC. Therefore this practice of studying a single scale at a time is often adopted in the modelling literature as a first step. For all these reasons [3, 4] adopted the simplifying assumption that the normative encoding problem faced by a given set of grid cells at a single point on the dorosventral axis of MEC may be effectively described as a single-scale problem.

### 6 Hexagonal grids are robust to small changes in place cell scale

Referring to their Fig. 4A (copied here in our Fig. 4A), [7] claims that *“The existence of grid solutions even with DoS readout encodings is highly sensitive to parameters such as the target function receptive field width σ*_*E*_, *and small alterations result in the disappearance of grid cells regardless of grid score threshold*.*”* For example in their Fig. 4A [7] claims that 10cm and 12cm place cell scales can generate grids but 11cm cannot.

This result is at odds with the theory of [3, 4], explained above, which predicts that grid scale will vary smoothly with place scale, specified by the inverse of the radius of the annulus in the Fourier domain in Fig. 1C. Indeed Fig. 4A of [7] (copied as our Fig. 4A) is also hard to reconcile with Fig. 3b of [7] (copied as our 4C), which actually confirms the theoretical prediction of [3, 4] that grid scale varies smoothly with place scale.

We thus performed our own direct test of the robustness of grid emergence to small changes in place scale by training path integrator networks over a range of place cell scales (0.11-0.13m, chosen because this range appeared to be particularly sensitive in Fig. 4A). We found a similar percentage of grid cells at all place cell scales and grid cell thresholds (Fig. 4B). Grid maps from these networks maintained their hexagonal structure and gradually changed scale as place cell scale was varied (Fig. 4D). Thus the emergence of hexagonal grid cells is robust with respect to minor variations in place cell scale, as expected from theory. We notice a major difference between Fig. 4A and Fig. 4B is the absolute percentage of grid cells above any grid score. For example, in Fig. 4A, only 10% to 15% of cells have grid score higher than 0.3, and less than 5% have grid score higher than 0.8. In contrast, networks in [4] have greater than 50% of cells with a grid score greater than 0.3 and greater than 20% of cells with a grid score greater than 0.8 (Supplemental Fig.2A pink histogram in [4]). The results of Fig. 4B are intermediate between these two: about 30% of cells with a grid score greater than 0.3 about 10% of cells with a grid score greater than 0.8. The primary difference between our results in Fig. 4B and those of [4] is that, in order to train many models quickly, we used an order of magnitude higher learning rate and an order of magnitude smaller training time. This higher learning rate lead to a large fraction of “dead ReLUs” (hidden units which never fired), and consequently lowered the absolute fraction of hexagonal grid cells. The optimization issue of dead ReLUs at high learning rates is a well known problem. Nevertheless, the fact that we still obtain many hexagonal grid cells illustrates that the results of [4] are at least robust to order of magnitude combined changes in learning rate and training time.

Based on this consideration of the very low absolute fraction of grid cells at all grid thresholds found in Fig. 4A in the work of [7], relative to that of [4] or the new simulations in Fig. 4B, we arrive at the following interpretation: rather than demonstrating “extreme sensitivity” of the emergence of hexagonal grids to place cell scale, [7] is simply reporting in Fig. 4A statistical fluctuations in grid score distributions in networks that do not robustly exhibit hexagonal grid cells to begin with.

### 7 Gaussian place cells can generate periodic patterns

[7] claims in Section 5.2 that they have proven mathematically the conclusion that *“a Gaussian interaction cannot produce periodic patterns*.*”* By this they mean that training path-integrators with place cell targets with “*Gaussian tuning curves should not produce periodic patterns*.” (Section 5.2 of [7]).

However, the conclusion of this mathematical proof is inconsistent with simulations. Indeed [2] found periodic patterns with Gaussian place cell tuning curves (Fig. 5A). Adapting their code and running it using a wider Gaussian place cell width (for theoretical reasons to be explained below), we found clear evidence of periodic patterns (stripes and squares) with Gaussian place cell targets (Fig. 5B).

We see two sources of incompleteness in the proof that explain this discrepancy with simulation results. First, the proof performs a linear stability analysis of a pattern formation problem with Gaussian interactions starting from a state of no patterns, as one increases the strength of the Gaussian interaction. In [7] they show that the first mode to become unstable is a zero frequency spatially homogeneous mode. This statement is correct. However, the conclusion that no periodic pattern can then emerge does not follow from this. As one increases the Gaussian interaction strength further, more modes with periodic patterns will also become unstable from a state of no patterns to begin with. These unstable periodic patterns may interact nonlinearly with the first unstable DC mode to generate an overall stable periodic pattern. Without showing that this does *not* happen, one cannot conclude that *“a Gaussian interaction cannot produce periodic patterns*.*”* For example, certain pattern formation problems with unimodal quadratic interactions and a quartic nonlinearity can favor square patterns.

A second source of incompleteness in the proof of [3] is that the pattern formation theory only addresses the ground state of a single optimization problem involving a single grid cell. When training a network to output place cell firing rates, there are multiple hidden units that could develop into grid cells each exhibiting slightly different patterns. Thus the global minimum of the pattern formation problem does not uniquely specify what every learned grid cell will do (it is likely that low energy configurations of the pattern formation problem at some finite temperature more correctly characterize the distribution of grid cells that appear in neural network training though we did not explore this). Nevertheless, with a wide Gaussian place cell profile, in Fourier space the DC mode (at the center of Fig. 1C) will have power, but also the 4 lowest frequency cardinal modes (cosines and sines in x and y, i.e. the 4 nearest lattice points to the center in Fig. 1C) will also have power, and a trained neural network will always do better in outputting place cells if it uses not only the DC mode but also the slightly higher frequency modes available to it in the output. The pure pattern formation analysis cannot account for these details.

But it is precisely this logic that lead us to consider training path-integrators with wide Gaussian place cell targets so that in Fourier space only the DC mode and the 4 cardinal lowest frequency modes had substantial power. We also used a tanh nonlinearity. Our results in Fig. 5B clearly show, consistent with theoretical considerations above, that our learned grid cells are dominated by random linear combinations of these 5 modes (DC, and sine and cosine in x and y). This often yields stripes and whenever there is a non-stripe lattice structure, the result is a square grid.

In summary, the conclusion of [7] that *“a Gaussian interaction cannot produce periodic patterns*,*”* is incorrect. Moreover this conclusion does not follow from the proof given in [7] because that proof is incomplete.

### 8 Biological plausibility of DoG inputs from grid to place cells

Finally, the authors of [7] point out that in experiments, biological place cells have mostly unimodal correlation structure, or at least lack an inhibitory surround, in contrast to the DoS receptive fields studied in [3, 4] as well as [14]. However this claim of biological implausibility made in [7] rests on an interpretation of place field training targets that is inconsistent with [4].

To clarify, we quote from [4]:

> *In the recurrent models, this surround corresponds to the layer of grid-like cells exciting the place cells with spatial RFs closest to the current position and inhibiting the neighboring place cells*.

Under this interpretation in [4], the center surround excitatory/inhibitory structure refers to *inputs* to place cells from the entire grid cell population *before* they are rectified to produce output place cell firing rates. Basically, the interpretation is *p_i_*(*x*) denotes the *input* current to place cells from the combined grid cell population when the animal is at location *x*. Such inhibition could lead to voltage responses in place cells that are subthreshold. A center-surround structure in the intepretation of [4] then means that this input current pattern across place cells should be anti correlated with itself for nearby location pairs.

This input current pattern is distinct from place cell output firing rates which both rectify these inputs and could receive inputs from additional cells other than grid cells that might potentially balance the subthreshold inhibition from grid cells. Because of the existence of rectifying nonlinearities and potentially additional inputs to place cells, correlations among superthreshold place cell firing rates may well be a poor predictor of correlations between sub-threshold place cell input currents specifically from the grid cell population. For example, assuming place cell output firing rates are rectified, an inhibitory surround input from grid cells could be totally invisible at the level of place cell firing rates, especially if this input were balanced by other inputs. For this reason, available experimental evidence for one sub-threshold correlation structure or another is weak, and so the DoG/DoS patterns of place cell input currents modeled in [3, 4] are difficult to rule out from considerations of place cell output firing rates alone.

## 9 Summary

In summary, we find, in contrast to claims made in [7], that hexagonal grid cells, under conditions delineated in [3, 4], do indeed: (1) robustly emerge under DoG place cell input patterns; (2) robustly emerge under place cells with multiple fields at a single scale; and (3) are robust to changes in place cell scale. Additionally we find, in contrast to [7], that Gaussian place cells can generate periodic patterns. Finally, we explain how DoG place cell input patterns from grid cells cannot easily be ruled out given existing experimental data.

Overall we find the results of [7], when critically assessed, are consistent with prior theoretical understanding of when hexagonal grid cells appear or not in trained path-integrator circuits.

